# Intercellular BVOC accumulation reflects sustained antioxidant defenses without additional carbon loss under ozone exposure in *Eugenia uniflora*

**DOI:** 10.64898/2026.08.10.743946

**Authors:** Alex do Nascimento, Fernanda Anselmo-Moreira, Bruno Ruiz Brandão da Costa, Matheus Casarini Siqueira, Claudia Maria Furlan, Silvia Ribeiro de Souza

## Abstract

Tropospheric ozone (O₃) is a major atmospheric pollutant that affects plant carbon metabolism, redox homeostasis, and secondary metabolism, including the biosynthesis and emission of biogenic volatile organic compounds (BVOCs). However, the contribution of BVOCs to O_3_ tolerance, particularly in tropical woody species, remains poorly understood. Here, we investigated whether acute O₃ exposure (cumulative AOT40 of 3497.82 ppb h) induces alterations in photosynthetic performance, redox homeostasis, and BVOC partitioning in *Eugenia uniflora*. We evaluated gas exchange, photosynthetic pigments, ascorbate and glutathione pools, emitted BVOCs, modeled intercellular BVOC concentrations, and the relative carbon cost associated with BVOC emissions. O₃ exposure significantly increased net CO₂ assimilation without affecting stomatal conductance, transpiration, leaf water status, or chlorophyll concentrations, indicating maintenance of photosynthetic performance. Carotenoid concentrations and total glutathione decreased, whereas glutathione redox status was maintained. O₃ induced marked compound-specific changes in BVOC composition and partitioning. Several monoterpenes appeared exclusively under O₃ exposure, γ-elemene emission increased significantly, and the relative distribution of individual BVOCs between the modeled intercellular and emitted pools was altered. These findings show that the response of *E. uniflora* to acute O₃ exposure was characterized by interplay among carbon assimilation, glutathione redox regulation, and BVOC partitioning rather than by increased total volatile emission. Enhanced carbon assimilation occurred without additional carbon loss through BVOC release, while changes in the modeled intercellular pool indicate that part of the volatile response remained within the leaf. Our findings highlight BVOC partitioning as an important dimension of the plant response to oxidative stress and demonstrate that emission measurements alone may not fully capture the fate and potential physiological role of volatile carbon under O₃ exposure.

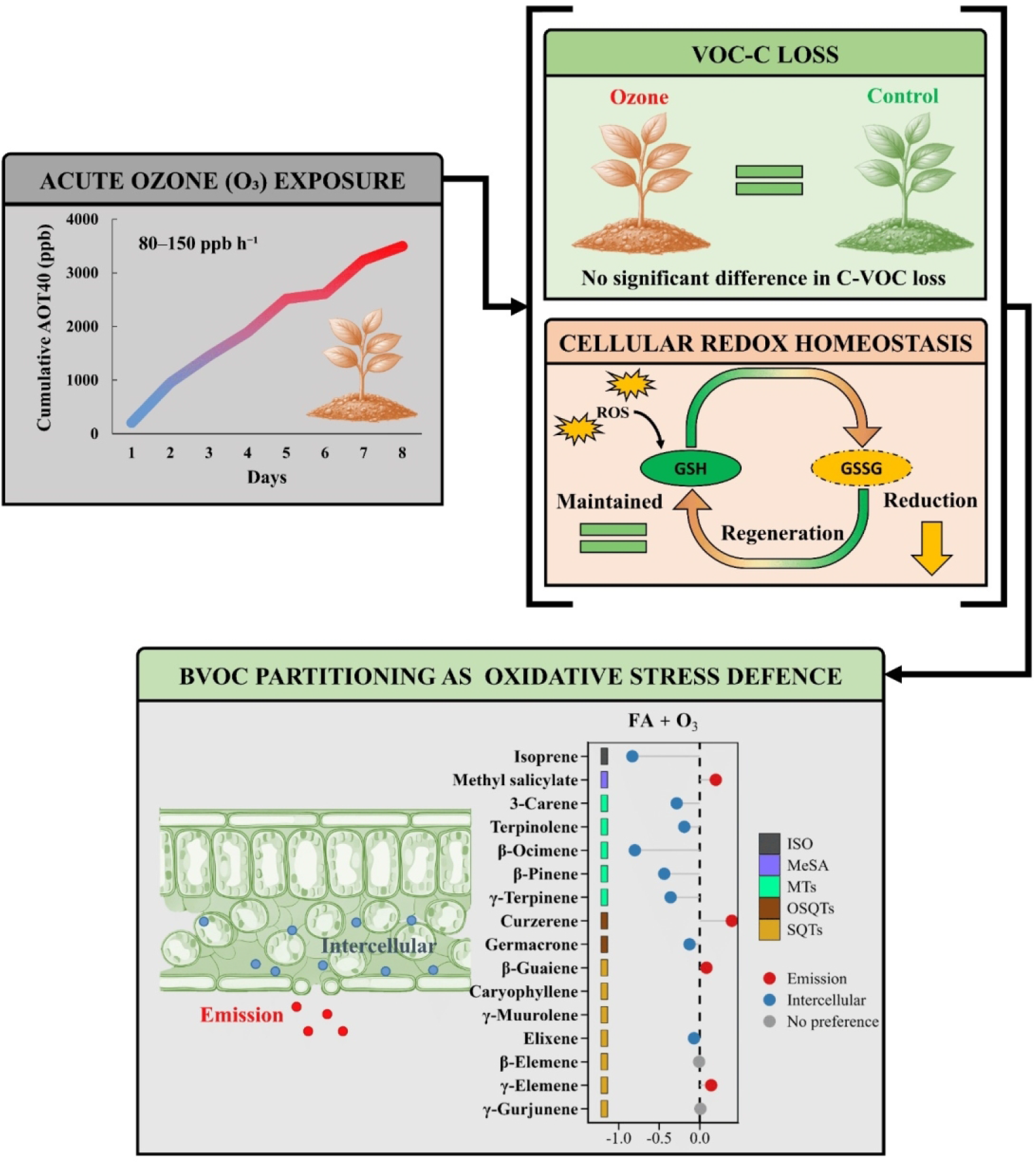

BVOC Partitioning Contributes to Oxidative Stress Defence Under Acute O₃ Exposure

## Introduction

Tropospheric ozone (O₃) is a photochemical pollutant formed through reactions involving nitrogen oxides and volatile organic compounds and is widely recognized as one of the most important atmospheric phytotoxic gases, affecting the vegetation across urban, peri-urban, agricultural, and natural environments (Mills et al. 2018, Borbon et al. 2026). O₃ can alter plant physiological by altering carbon assimilation, redox homeostasis, and specialized metabolism, often before visible injury becomes apparent. After entering leaves mainly through stomata, O₃ reacts rapidly in the apoplast and promotes the formation of reactive oxygen species (ROS), which can trigger oxidative imbalance, metabolic alterations, and, depending on exposure intensity and plant sensitivity, reductions in photosynthesis, growth, and biomass accumulation (Ainsworth et al. 2012; Vainonen and Kangasjärvi 2015).

Plant responses to O₃-induced oxidative stress depend on the coordination between stomatal regulation, photosynthetic metabolism, and antioxidant capacity. Stomatal closure may reduce pollutant uptake, but it can also restrict CO₂ diffusion and limit carbon assimilation, creating a trade-off between protection and carbon gain under polluted atmospheres (Ainsworth et al. 2012; Mills et al. 2018). Once oxidative stress is established, plants activate mechanisms to maintain cellular redox homeostasis, reflected by the redox potential of the ascorbate and glutathione pools, commonly assessed through the ratios of reduced ascorbate to dehydroascorbate (AsA/DHA) and reduced to oxidized glutathione (GSH/GSSG), together with antioxidant metabolites such as carotenoids (Gill and Tuteja 2010; Foyer and Noctor 2011; Noctor et al. 2012).

Beyond classical antioxidant systems, biogenic volatile organic compounds (BVOCs) represent an additional, often underrecognized component of plant defense against abiotic stress (Zuo et al. 2025). These compounds are not only products released to the atmosphere, but also metabolites with potential physiological functions inside leaves, particularly when oxidative stress increases (Peñuelas and Staudt 2010; Zuo et al. 2025). Among them, volatile isoprenoids such as isoprene, monoterpenes, and sesquiterpenes have been associated with protection against oxidative damage through reactions with oxidants, membrane stabilization, reduced lipid peroxidation, and modulation of redox signalling (Costa et al. 2025, Zuo et al. 2025). This protective function may occur before BVOCs are emitted to the atmosphere, because volatile compounds can accumulate or partition within membranes, cell walls, and leaf internal air spaces, where they may interact with oxidants locally (Niinemets et al. 2014). However, because most studies quantify only emitted BVOCs, the dynamics and physiological significance of intercellular volatile pools remain largely unexplored, despite their potential role in antioxidant defense and carbon investment during oxidative stress. In addition, BVOC production represents a carbon investment that may constitute a substantial fraction of recently assimilated carbon under O_3_ stress, suggesting a potential trade-off between primary metabolism and volatile-mediated defense. Recent studies further indicate that the intercellular air space is an active site of O_3_ defense, where the balance between stomatal conductance and intercellular BVOC concentrations determines the potential for local O_3_ scavenging before oxidants reach living tissues (Yu and Blande 2021; Yu and Blande 2022).

Despite increasing evidence that BVOCs contribute to antioxidant defense, it remains unclear whether their metabolism constitutes an active acclimation strategy supported by carbon investment or merely a secondary consequence of oxidative stress. Consequently, the integration between carbon assimilation, intercellular BVOC, and cellular redox regulation during O_3_ stress remains poorly understood. This knowledge gap is particularly relevant for tropical woody species, which rely on chemically diverse specialized metabolism to cope with oxidative stress but remain much less studied than temperate trees or crop species in terms of integrated physiological, biochemical, and volatile responses (Mills et al. 2018; Moura et al. 2022; Cheesman et al. 2024).

Among tropical woody species, *Eugenia uniflora* L. (Myrtaceae), commonly known as pitangueira or Surinam cherry, is a neotropical tree native to South America and widely distributed in Brazil, where it occurs in natural, urban, peri-urban, and restoration settings (Engela et al. 2021), provides an attractive model. Its frequent use in urban landscaping, degraded-area restoration, and domestic orchards gives the species relevance for studies of vegetation responses to environmental pollution (Anselmo-Moreira et al. 2025). In addition, terpene-rich volatile profiles, often with a substantial contribution of sesquiterpenes, have been reported for several native Atlantic Forest species (Costa et al. 2020; Anselmo-Moreira et al. 2025), including *E. uniflora* (Anselmo-Moreira et al. 2026), supporting its suitability for investigating stress-related BVOC responses. Previous work has also shown that *E. uniflora* is sensitive to chronic O₃ exposure, exhibiting physiological and metabolic alterations consistent with oxidative stress (Engela et al. 2021). Moreover, these studies indicate that the response of *E. uniflora* to O₃ is strongly dependent on the exposure regime. Chronic exposure to elevated O₃ has been shown to induce substantial physiological and metabolic alterations, including reductions in photosynthetic performance and changes in antioxidant defenses and primary metabolism, even in the absence of pronounced visible foliar injury (Engela et al. 2021). In contrast, acute O₃ exposure can trigger antioxidant depletion and selective shift in BVOC and metabolomic profiles while major physiological functions remain comparatively stable (Anselmo-Moreira et al. 2025). Previous observations also suggest that changes in BVOC emission may not necessarily reflect changes in the internal volatile pool, raising the possibility that O₃ alters the partitioning of BVOCs between their accumulation within the leaf and their release to the atmosphere. This distinction is particularly relevant because internally retained BVOCs may contribute to antioxidant protection, membrane stabilization, or reactions with incoming O₃ before these compounds are emitted.

Thus, *E. uniflora* provides an excellent system for investigating the integration of carbon assimilation, redox regulation, and BVOC partitioning under acute oxidative stress. Here, we hypothesized that acute O₃ exposure triggers a reorganization of BVOC partitioning, altering the balance between intercellular retention and atmospheric emission while promoting coupled changes in carbon assimilation and redox homeostasis. This framework considers BVOC metabolism as an actively regulated, rather than solely constitutive, component of plant acclimation to oxidative stress. Understanding these mechanisms may provide new insights into how BVOC partitioning contributes to O_3_ tolerance by linking carbon investment with cellular redox homeostasis in tropical woody species.

## Material and methods

### Plant material and ozone exposure

Juvenile plants of *E. uniflora* were commercially obtained from a commercial plant nursery (Fábrica de Florestas, Bragança Paulista, São Paulo, Brazil) and transplanted into 3 L pots containing Carolina Soil substrate. At the time of the experiment, the saplings were approximately 50 cm tall. Before O_3_ exposure, plants were maintained for four weeks in a greenhouse under controlled temperature conditions (25–30°C). During this acclimation period, plants were irrigated daily and fertilized weekly with 250 mL of Hoagland nutrient solution.

One day before the beginning of the fumigation experiment, all plants were irrigated to field capacity. After water drainage, the pots were wrapped in plastic bags to reduce water loss from the substrate. O_3_ exposure experiments were conducted at the Laboratory of Plant-Atmospheric Interaction (LABIAP), of the Environmental Research Institute of São Paulo (IPA-SP), Brazil. The experiment was carried out in closed chambers made of stainless steel and lined with Teflon (85 × 94 × 85 cm), as described by Souza and Pagliuso (2009). The chambers were maintained under controlled temperature and artificial lighting conditions. Light was supplied by full-spectrum LED panels developed for indoor plant cultivation, providing a photosynthetic photon flux density of approximately 700 µmol m⁻² s⁻¹. The photoperiod was set from 08:00 to 16:30.

Plants were divided into two experimental groups: filtered air (FA) and filtered air enriched with O_3_ (FA + O₃), with nine plants per treatment. The air supplied to the chambers was purified with an efficiency of 98.5%. In the O_3_ treatment, FA was enriched with O₃ concentrations ranging from 80 to 150 ppb, which were continuously monitored throughout the exposure period (Figure S1 and Table S1 available as Supplementary Data at Tree Physiology Online). O_3_ fumigation was performed for 5 h per day, from 09:00 to 14:00, over eight consecutive days, resulting in acute ozone exposure, which consists of a short-term fumigation that rapidly reaches a critical accumulated dose (AOT40 of 3497.82 ppb·h). Plant positions within the chambers were randomized daily to minimize positional effects.

### Physiological parameters

#### Leaf pH

For each plant, leaf pH was determined by macerating 1.5 g of fresh leaf tissue in 15 mL of deionized water. Ten milliliters of the supernatant were collected using a plastic syringe and transferred to a 15 mL beaker. The pH was measured using a portable multiparameter meter (Combo 5, AKSO, Brazil). (Pérez-Harguindeguy et al 2013, Liu et al. 2022)

#### Relative Water Content (RWC)

RWC was determined for each plant using five leaf discs (0.7 cm in diameter), collected from each plant. The discs were immediately weighed to obtain fresh mass (FM), hydrated in 25 mL of distilled water for 24 h under refrigeration in the dark, and then weighed to obtain turgid mass (TM). Subsequently, the discs were oven-dried at 60 °C for 72 h to determine dry mass (DM). RWC was calculated according to González and González-Vilar (2001) and Anselmo-Moreira et al. (2026) using the following equation:

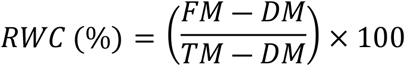

#### Leaf Gas Exchange

Leaf gas exchange measurements were performed simultaneously with BVOC sampling using a CO₂/H₂O gas analyzer system (LI-850, LI-COR Inc., Lincoln, NE, USA). The analyzer continuously monitored CO₂ and H₂O concentrations inside the Teflon sampling bags during the enclosure period. Transpiration rate (*E*), CO₂ assimilation rate (*A*), and stomatal conductance to water vapor (*g*_sw_) were estimated following the principles described by von Caemmerer and Farquhar (1981). Additional details on the gas exchange calculations are provided in the Supplementary Data at Tree Physiology Online.

#### Photosynthetic Pigments

Photosynthetic pigments were quantified from 50 mg of ground leaf tissue extracted in 10 mL of 95% ethanol. Samples were vortexed for 10 s and kept in the dark at 4 °C for 24 h. After extraction, the supernatant was transferred to quartz cuvettes, and absorbance was measured at 470, 649, and 664 nm using a UV–Vis spectrophotometer (Genesys 10S, Thermo Fisher Scientific, USA). Chlorophyll *a*, chlorophyll *b*, total chlorophyll, and total carotenoids (Car) were calculated according to the equations described by Lichtenthaler (1987) and Minocha et al. (2009), and results were expressed as μg g⁻¹ fresh mass.

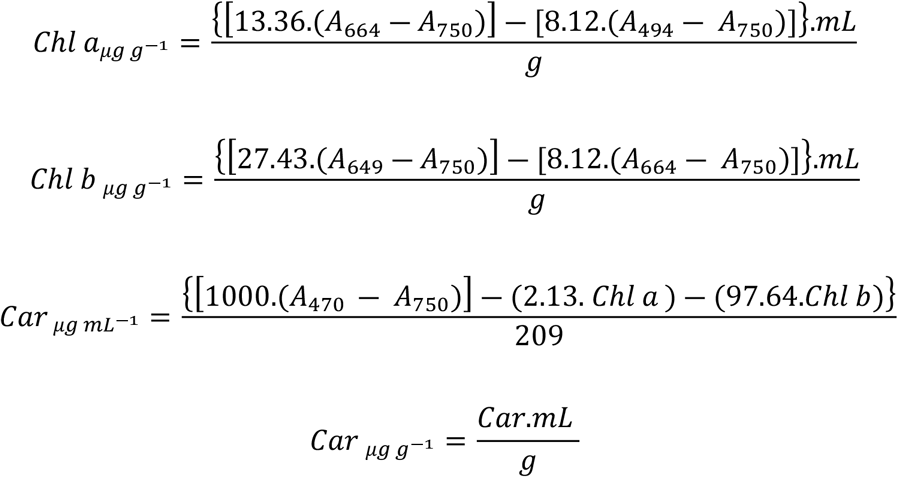

### Redox status: ascorbate and glutathione pools

Reduced and total pools of ascorbate and glutathione were determined by high-performance liquid chromatography coupled to a diode-array detector (HPLC-DAD; 1260 Infinity, Agilent Technologies, USA). Chromatographic separation was performed using an Eclipse XDB-C18 column (150 × 4.6 mm, 5 µm) maintained at 25 °C. The mobile phase consisted of ultrapure water acidified with phosphoric acid (H₃PO₄, pH 2.3), delivered under isocratic conditions at a flow rate of 1 mL min⁻¹ and run time of 20 min. Reduced ascorbate (AsA) and reduced glutathione (GSH) were monitored at 245 and 194 nm, respectively.

Frozen and ground leaf tissue (150 mg) was homogenized in 2 mL of 6% metaphosphoric acid (HPO₃) containing 1 mM ethylenediaminetetraacetic acid disodium salt dihydrate (EDTA-Na₂). After centrifugation at 10,000 rpm for 15 min at 4 °C, the supernatant was used for chromatographic analysis. Reduced forms were determined by diluting 100 µL of supernatant with 400 µL of mobile phase, followed by filtration (PTFE 0.45 µm), and 20 µL was injected into the HPLC system. Total ascorbate and total glutathione were determined by mixing 100 µL of supernatant with 20 µL of 0.4% dithiothreitol (DTT), prepared in 2 M sodium phosphate buffer (pH 7.0), and 10 µL of 45% K₂HPO₄. The mixture was incubated for 20 min in the dark on ice, and the reaction was stopped by adding 20 µL of 2 M H₃PO₄. The final volume was adjusted with 350 µL of ultrapure water, filtered (PTFE 0.45 µm), and 20 µL were injected into the HPLC (López et al. 2005; Sala-Carvalho et al. 2022).

Calibration curves were constructed with the intercept fixed at zero, assuming a null UV detector response in the absence of analyte and based on the linear distribution of the experimental data supporting this assumption. A total of 18 concentration points (0.5–26 µg mL⁻¹) were used, resulting in the following regression equations: for ascorbic acid, y = 0.4112·x (R² = 0.9983); and for glutathione, y = 0.4241·x (R² = 0.9979). Dehydroascorbate (DHA) and oxidized glutathione (GSSG) were estimated as the difference between the total and reduced pools. Redox status was expressed as the ratios AsA/total ascorbate and GSH/total glutathione. Results were expressed as µg g⁻¹ fresh mass.

### BVOC sampling and analysis

BVOCs were collected immediately after O_3_ exposure from plants of each treatment. For each plant, branches were enclosed in Teflon bags under a continuous flow of clean air (2.5 ± 0.5 L min⁻¹). Incoming air was purified through a filtration system containing glass wool for particle removal, silica gel, activated charcoal, and potassium permanganate. Emitted VOCs were trapped on cartridges containing 100 mg of Tenax TA (60/80 mesh), connected to suction pumps (Air Lite, SKC) operating at 250 mL min⁻¹ for 1 h (Figure. S2). After sampling, enclosed leaves were scanned to determine leaf area using a leaf area meter (LI-3100C, LI-COR Inc., Lincoln, NE, USA). Leaves were then oven-dried at 60 °C for 72 h and weighed to obtain dry mass. After collection, cartridges were stored in a desiccator with silica gel until analysis.

BVOC analysis was performed using gas chromatography–mass spectrometry (GC–MS; Agilent 7890B, 5977A). Thermal desorption was carried out using an automated thermal desorption system (ATD 650, PerkinElmer) under nitrogen flow at 250 °C for 10 min. Analytes were cryofocused at −30 °C prior to injection and separated on an HP-5 capillary column (50 m × 0.2 mm × 0.5 µm). Helium was used as the carrier gas. The oven temperature program was as follows: 46 °C for 5 min, increased at 5 °C min⁻¹ to 210 °C and held for 20 min, followed by an increase at 10 °C min⁻¹ to 250 °C and held for 5 min. Mass spectra were acquired in scan mode over an *m/z* range of 50–750

Compound annotation was performed by comparing mass spectra with the NIST library (version 2.0g, 2012) with matches higher than 800. Quantification was based on external calibration curves prepared with commercial standards (Sigma-Aldrich). When compound-specific standards were unavailable, a representative standard from the same chemical class was used for estimation of quantification. The calibration curve of 3-carene was applied to isoprene (ISO) and monoterpenes (MTs), methyl salicylate to methyl salicylate (MeSA), β-caryophyllene to non-oxygenated sesquiterpenes (SQTs), and farnesol to oxygenated sesquiterpenes (OSQTs) (Table S2 avaliable as Supplementary Data at Tree Physiology Online)

BVOC emission rates (*ER*, ng gDM⁻¹ h⁻¹) were calculated from the sampled air concentration, total sampled air volume, sampling duration, and dry mass of the enclosed leaves:

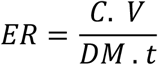

where *C* is the sampled air concentration (ng L⁻¹), *V* is the sampled air volume (L), *DM* is the dry mass of the enclosed leaves (g), and *t* is the sampling duration (h).

Emission rates were then normalized (*ER*_n_, ng gDM⁻¹ h⁻¹) to a reference temperature of 303 K using a temperature correction algorithm adapted from Guenther et al. (2012): where *T* is the temperature inside the sampling bag, *T*_s_ is the standard temperature (set to 303 K), and β is an empirical coefficient. A β value of 0.17 was used for sesquiterpenes (SQTs and OSQTs), whereas 0.10 was used for MTs and other VOCs. Emission rates were converted from ng gDM⁻¹ h⁻¹ to µg gDM⁻¹ h⁻¹ and are reported in these units throughout the manuscript for consistency.

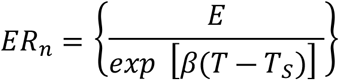

### Model-based Estimation of Intercellular BVOC Concentrations

Intercellular BVOC concentrations were estimated using a model that assumes that BVOC emissions from foliage occur predominantly through stomatal diffusion. This approach was based on classical diffusion models described by Fall and Monson (1992) and Niinemets et al. (2002), as applied by Yu and Blande (2022). BVOC emission can be described as:

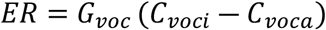

where *ER* is the BVOC emission rate, C_voci_ and C_voca_ correspond to the intercellular and ambient BVOC concentrations, respectively, both expressed in mol mol⁻¹. G_voc_ represents the total conductance for a given BVOC, including stomatal and boundary layer conductance, expressed in mol m⁻² s⁻¹. The equation was rearranged to estimate intercellular BVOC concentration:

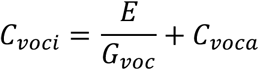

G_voc_ was calculated from stomatal conductance (G_svoc_) and boundary layer conductance (G_bvoc_) for each BVOC, as follows:

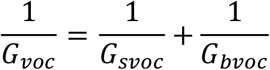

Compound-specific G_svoc_ and G_bvoc_ were estimated according to Burrows and Milthorpe (1976) as follows:

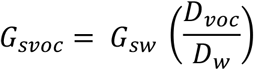

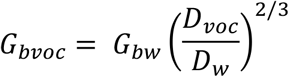

where *g*_sw_ is stomatal conductance to water vapor and G_bw_ is boundary layer conductance to water vapor, both expressed in mol m⁻² s⁻¹. A fixed value of 2.5 mol m⁻² s⁻¹ was used for G_bw_, following LI-COR Inc. (2002).

The ratio between the gas-phase diffusion coefficient of each BVOC (D_voc_) and that of water vapor (D_w_) was estimated from their molecular masses:

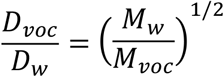

Where M_w_ and M_voc_ are the molecular masses of water (18 g mol⁻¹) and each BVOC, respectively, expressed in g mol⁻¹.

### Estimation of Carbon Loss Fraction via BVOCs

To estimate the fraction of carbon lost through BVOC emissions, emission rates were first expressed on a leaf-area basis:

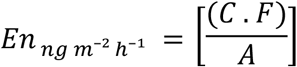

where *En* is the area-based BVOC emission rate (ng m⁻² h⁻¹), *C* is the sampled BVOC concentration (ng L⁻¹), *F* is the inlet airflow rate (L h⁻¹), and *A* is the enclosed leaf area (m²).

The proportion of carbon lost via BVOC emissions was determined according to Yu and Blande (2021):

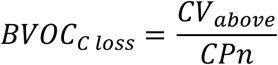

where *CV_above_* is the carbon-based BVOC flux above the leaf surface (ng C m⁻² h⁻¹) and *CP*n represents the carbon assimilated through net photosynthesis by aboveground tissues, expressed as ng C m⁻² h⁻¹. *CV_above_* was calculated as:

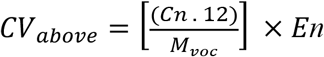

where *Cn* is the number of carbon atoms specific to each BVOC. For presentation purposes, carbon fluxes were converted from ng Cm⁻² h⁻¹ to mg Cm⁻² h⁻¹ and are reported in these units throughout the manuscript.

### Data Analysis

The relative contribution of each BVOC to the total detected profile was calculated separately for emitted and intercellular compounds in each treatment.

The total carbon loss via BVOC emissions was calculated for each individual as the sum of the carbon losses associated with all quantified BVOCs, expressed in mg C m⁻² h⁻¹. Relative carbon loss (%) was then calculated as the ratio between total BVOC-derived carbon loss and the corresponding carbon assimilated through net photosynthesis (CPn), multiplied by 100. For statistical analyses based on proportional data, percentage values were converted to proportions by dividing by 100.

All statistical analyses were performed in R software (R Core Team, 2024). Physiological and biochemical parameters were compared between treatments (FA and FA + O₃), using Student’s t-test when assumptions of normality were met. Data normality was assessed using the Shapiro–Wilk test, and appropriate transformations were applied when necessary. When normality was not achieved, the Wilcoxon rank-sum test was used. For emitted and modeled intercellular BVOCs, univariate comparisons between treatments were performed only for compounds detected in both groups. Differences were assessed using the Wilcoxon rank-sum test, and effect size was estimated using the rank-biserial correlation coefficient (*r*) to evaluate the magnitude and direction of O_3_ effects. Absolute *r* values were interpreted as small (∼0.10), moderate (∼0.30), or large (≥0.50) effects Cohen (1988). Positive *r* values indicated higher values in O_3_-exposed plants, whereas negative values indicated higher values in controls. Effect sizes were calculated using the effectsize package. Differences between treatments were evaluated using either parametric or non-parametric tests, depending on data distribution and variance homogeneity.

For multivariate analyses of BVOC profiles, compound retention and imputation were assessed separately within each treatment. BVOCs detected in at least 40% of individuals within a treatment were retained, and remaining missing values were imputed using missForest (Stekhoven and Bühlmann 2012), with iterations stopped when the change in out-of-bag error was < 0.001. BVOCs detected in fewer than 40% of individuals within a treatment were considered absent in that treatment and were not imputed. A pseudocount of 1 × 10⁻⁴ was added to zero values before multivariate analyses.

Principal component analysis (PCA) was performed on centered and scaled BVOC emission data using the prcomp function in R. PCA scores and loadings were extracted to evaluate treatment-related patterns and identify compounds contributing to group separation. PCA results were visualized using the FactoMineR (Lê and Husson 2008) and factoextra packages (Kassambara and Mundt 2020), with individuals projected onto the first two principal components and grouped according to treatment. Ninety-five percent confidence ellipses were added to represent group dispersion, and BVOC variables were displayed in biplots according to their contribution to the ordination. To statistically assess treatment effects on BVOC composition, data were transformed using the centered log-ratio (CLR) transformation (Filzmoser et al. 2018). Permutational analysis of variance (PERMANOVA) was performed using the adonis2 function from the vegan package, with Euclidean distance and 999 permutations. Homogeneity of multivariate dispersion was evaluated using betadisper, followed by analysis of variance.

To evaluate the relative distribution of BVOCs between the modeled intercellular and emitted pools, intracellular concentration data and emission rates were initially transformed into relative proportions. This normalization was performed individually for each sample by considering the percentage contribution of each compound relative to the total detected BVOCs, allowing the comparison of relative composition between the two pools independently of differences in absolute magnitudes.

The relative partitioning of each BVOC was estimated by calculating the base-2 logarithm of the ratio between the emitted fraction and the intracellular fraction.Positive LR values indicated a greater relative representation of the compound in the emitted fraction, whereas negative values indicated greater relative representation in the intercellular pool. Values close to zero were interpreted as indicating no preferential partitioning between the two pools. For each combination of treatment (FA and FA + O₃) and individual compound, the median LR values were calculated. The hypothesis that relative partitioning differed from a distribution centered at zero was tested using a one-sample Wilcoxon signed-rank test, considering LR = 0 as the expected value under the absence of preferential partitioning.

## Results

### Physiological and biochemical responses

Acute O₃ exposure induced selective physiological and biochemical alterations in *E. uniflora* seedlings (Table 1). Among the gas-exchange variables, *A* showed the strongest response, increasing by approximately 135% in O₃-exposed plants relative to seedlings maintained under filtered air (FA). In contrast, *g*_sw_, ER, and (RWC remained unchanged between treatments, despite a tendency toward higher mean *g*_sw_ values under ozone exposure (FA+O₃).

**Table 1.** Effects of ozone (O₃) exposure on physiological and biochemical in *Eugenia uniflora* seedlings.

| Student's t-test |  |  |  |  |  |  |  |
| --- | --- | --- | --- | --- | --- | --- | --- |
| Parameter | Treatment | Mean | SD | Range (min–max) | Test statistic | *p* value | Significant |
| A (μmol m<sup>-2</sup> s<sup>-1</sup>) | FA | 3.25 | 1.48 | 1.05-5.84 | -3.19 | 0.009 | Yes |
|  | FA + O<sub>3</sub> | 7.64 | 3.85 | 1.89-11.87 |  |  |  |
| g<sub>sw</sub> (mol m<sup>-2</sup> s<sup>-1</sup>) | FA | 0.09 | 0.04 | 0.02-0.13 | -2.09 | 0.069 | No |
|  | FA + O<sub>3</sub> | 0.27 | 0.26 | 0.03-0.81 |  |  |  |
| E (mmol m<sup>-2</sup> s<sup>-1</sup>) | FA | 1.36 | 0.42 | 0.53-1.77 | -0.41 | 0.691 | No |
|  | FA + O<sub>3</sub> | 1.47 | 0.66 | 0.65-2.62 |  |  |  |
| Carotenoids (mg g<sup>-1</sup>) | FA | 0.16 | 0.04 | 0.10-0.23 | 2.86 | 0.015 | Yes |
|  | FA + O<sub>3</sub> | 0.11 | 0.02 | 0.08-0.14 |  |  |  |
| AsA (μg mg<sup>-1</sup>) | FA | 0.27 | 0.05 | 0.20-0.35 | 1.73 | 0.111 | No |
|  | FA + O<sub>3</sub> | 0.2 | 0.11 | 0.04-0.31 |  |  |  |
| DHA (μg mg<sup>-1</sup>) | FA | 0.89 | 0.43 | 0.42-1.77 | -0.94 | 0.365 | No |
|  | FA + O<sub>3</sub> | 1.12 | 0.61 | 0.43-2.00 |  |  |  |
| Total AsA (μg mg<sup>-1</sup>) | FA | 1.16 | 0.4 | 0.77-1.97 | -0.65 | 0.529 | No |
|  | FA + O<sub>3</sub> | 1.32 | 0.64 | 0.66-2.21 |  |  |  |
| GSSG (μg mg<sup>-1</sup>) | FA | 1.48 | 0.26 | 1.07-1.88 | 2.85 | 0.013 | Yes |
|  | FA + O<sub>3</sub> | 1.05 | 0.37 | 0.40-1.56 |  |  |  |
| Total GSH (μg mg<sup>-1</sup>) | FA | 1.66 | 0.26 | 1.27-2.10 | 2.79 | 0.015 | Yes |
|  | FA + O<sub>3</sub> | 1.22 | 0.4 | 0.55-1.87 |  |  |  |
| Wilcoxon rank-sum test |  |  |  |  |  |  |  |
| Parameter | Treatment | Median | Q1 | Q3 | Test Statistic | *p* value | Significant |
| pH | FA | 5.08 | 5.01 | 5.16 | 65 | 0.034 | Yes |
|  | FA + O<sub>3</sub> | 4.86 | 4.85 | 4.95 |  |  |  |
| RWC (%) | FA | 59.25 | 52.3 | 65.59 | 36 | 0.73 | No |
|  | FA + O<sub>3</sub> | 62.36 | 60.2 | 65.2 |  |  |  |
| Chlorophyll *a* (mg g<sup>-1</sup>) | FA | 0.44 | 0.4 | 0.55 | 58 | 0.136 | No |
|  | FA + O<sub>3</sub> | 0.36 | 0.35 | 0.45 |  |  |  |
| Chlorophyll *b* (mg g<sup>-1</sup>) | FA | 0.13 | 0.12 | 0.18 | 37 | 0.796 | No |
|  | FA + O<sub>3</sub> | 0.13 | 0.13 | 0.18 |  |  |  |
| Total chlorophyll (mg g<sup>-1</sup>) | FA | 0.54 | 0.52 | 0.73 | 53 | 0.297 | No |
|  | FA + O<sub>3</sub> | 0.49 | 0.48 | 0.65 |  |  |  |
| GSH (μg mg<sup>-1</sup>) | FA | 0.15 | 0.13 | 0.22 | 40 | 1 | No |
|  | FA + O<sub>3</sub> | 0.17 | 0.15 | 0.18 |  |  |  |
| AsA/Total AsA | FA | 0.22 | 0.19 | 0.31 | 62 | 0.063 | No |
|  | FA + O<sub>3</sub> | 0.12 | 0.08 | 0.19 |  |  |  |
| GSH/Total GSH | FA | 0.1 | 0.08 | 0.15 | 27 | 0.258 | No |
|  | FA + O<sub>3</sub> | 0.16 | 0.1 | 0.16 |  |  |  |

Values are presented as mean, standard deviation (SD), and range for variables analyzed using Student’s t-test, and as median, first quartile (Q1), and third quartile (Q3) for variables analyzed using the Wilcoxon rank-sum test. The test statistic corresponds to t for Student’s t-test and W for the Wilcoxon rank-sum test (n = 9 per treatment). Differences were considered significant at p < 0.05. Abbreviations: FA, filtered air; FA + O<sub>3</sub>, ozone-exposed plants; A, net CO<sub>2</sub> assimilation rate (μmol m<sup>-2</sup> s<sup>-1</sup>); g<sub>sw</sub>, stomatal conductance to water vapor (mol m<sup>-2</sup> s<sup>-1</sup>); E, transpiration rate (mmol m<sup>-2</sup> s<sup>-1</sup>); RWC, relative water content (%); Chlorophyll a, b, Total and carotenoids (mg g<sup>-1</sup>); AsA, reduced ascorbate (μg mg<sup>-1</sup>); DHA, dehydroascorbate (μg mg<sup>-1</sup>); GSH, reduced glutathione (μg mg<sup>-1</sup>) and GSSG, oxidized glutathione (μg mg<sup>-1</sup>)

Despite the maintenance of leaf hydration, FA+O₃ treatment altered leaf chemical characteristics, reducing leaf pH from a median of 5.08 to 4.86 in O₃-treated plants (Table 1). Photosynthetic pigments exhibited a similarly selective response. Carotenoid concentration declined by approximately 31% following O₃ exposure, whereas chlorophyll *a*, chlorophyll *b*, and total chlorophyll remained unchanged, indicating FA+O₃ treatment affected carotenoid accumulation without detectable changes in the chlorophyll pool.

The antioxidant systems also responded differentially to FA+O₃ treatment. The ascorbate pool remained remarkably stable, with no significant changes in AsA, DHA, total ascorbate, or the AsA/total AsA ratio between treatments (Table 1). Although the AsA redox ratio tended to decline under FA+O₃ treatment, this trend was not statistically significant. In contrast, the glutathione pool exhibited a selective depletion. GSSG decreased by approximately 29%, accompanied by a 27% reduction in total glutathione, whereas GSH remained unchanged. Consequently, the GSH/total glutathione ratio was maintained despite the reduction in the overall glutathione pool (Table 1).

### Carbon assimilation and BVOC-derived carbon loss

Despite the marked increase in carbon uptake, carbon loss through BVOC emissions remained unchanged between treatments (Figure 1 a,b, Table S3 available as Supplementary Data at Tree Physiology Online). Likewise, the proportion of recently assimilated carbon released as BVOCs did not differ significantly, although O₃-treated plants consistently exhibited lower median values than filtered-air plants (Figure 1c). Thus, enhanced carbon assimilation under FA+O₃ treatment was not accompanied by a proportional increase in carbon loss through volatile emissions, indicating that the additional assimilated carbon remained available for metabolic processes other than volatile release.

**Figure 1.**
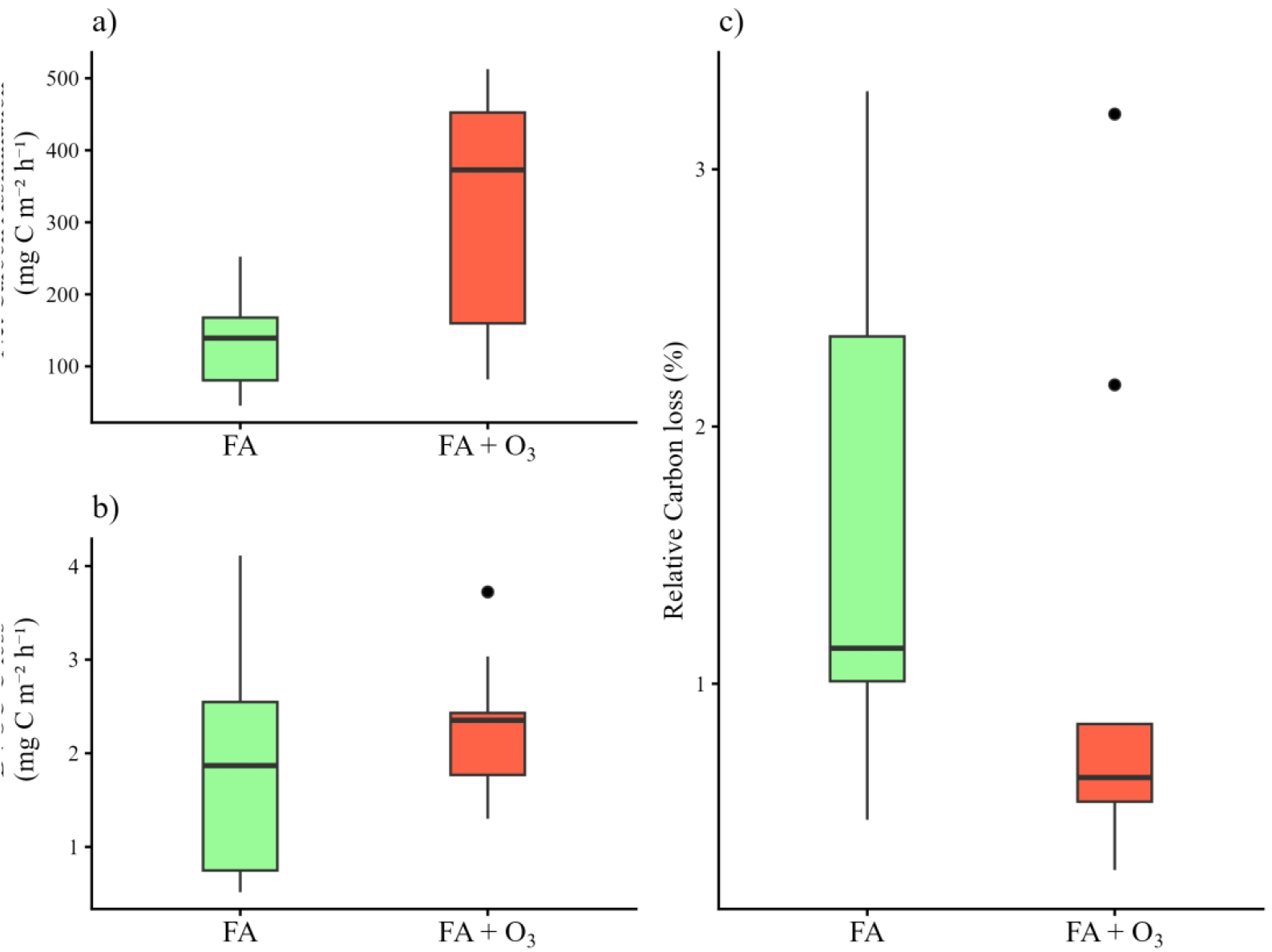
Tropospheric ozone (O₃) is a major atmospheric pollutant that affects plant carbon metabolism, redox homeostasis, and secondary metabolism, including the biosynthesis and emission of biogenic volatile organic compounds (BVOCs). However, the contribution of BVOCs to O3 tolerance, particularly in tropical woody species, remains poorly understood. Here, we investigated whether acute O₃ exposure (cumulative AOT40 of 3497.82 ppb h) induces alterations in photosynthetic performance, redox homeostasis, and BVOC partitioning in Eugenia uniflora. We evaluated gas exchange, photosynthetic pigments, ascorbate and glutathione pools, emitted BVOCs, modeled intercellular BVOC concentrations, and the relative carbon cost associated with BVOC emissions. O₃ exposure significantly increased net CO₂ assimilation without affecting stomatal conductance, transpiration, leaf water status, or chlorophyll concentrations, indicating maintenance of photosynthetic performance. Carotenoid concentrations and total glutathione decreased, whereas glutathione redox status was maintained. O₃ induced marked compound-specific changes in BVOC composition and partitioning. Several monoterpenes appeared exclusively under O₃ exposure, γ-elemene emission increased significantly, and the relative distribution of individual BVOCs between the modeled intercellular and emitted pools was altered. These findings show that the response of E. uniflora to acute O₃ exposure was characterized by interplay among carbon assimilation, glutathione redox regulation, and BVOC partitioning rather than by increased total volatile emission. Enhanced carbon assimilation occurred without additional carbon loss through BVOC release, while changes in the modeled intercellular pool indicate that part of the volatile response remained within the leaf. Our findings highlight BVOC partitioning as an important dimension of the plant response to oxidative stress and demonstrate that emission measurements alone may not fully capture the fate and potential physiological role of volatile carbon under O₃ exposure.

### BVOC emission rates and modeled intercellular concentrations

O₃ exposure markedly altered the composition of BVOC emissions in *E. uniflora* without increasing total BVOC-derived carbon loss. A total of 16 BVOCs were detected across treatments, including ISO, MeSA, MTs, OSQTs, and SQTs. SQTs and OSQTs were the dominant emitted classes under both treatments, although the relative contribution of individual compounds changed substantially in FA+O₃ treatment (Table 2). Also, this treatment altered both the qualitative and quantitative composition of the emission profile. Under FA treatment, the emission profile was mainly composed of elixene, germacrone, γ-elemene, curzerene, and caryophyllene. In addition, caryophyllene and γ-muurolene occurred only in this group. Under FA+O₃ treatment, six compounds were detected, including monoterpenes 3-carene, β-pinene, β-ocimene, γ-terpinene, and terpinolene together with the sesquiterpene β-guaiene. Among compounds detected in both treatments, γ-elemene showed the strongest positive response, with an emission rate approximately 2.5-fold higher under FA+O₃ treatment. In contrast, the emission rates of ISO, MeSA, curzerene, germacrone, elixene, β-elemene, and γ-gurjunene did not differ significantly between treatments. Notably, all detected MTs were induced by O₃ exposure, whereas ISO, MeSA, curzerene, germacrone, elixene, β-elemene, γ-elemene, and γ-gurjunene were present under both treatments and therefore represented constitutive compounds of the volatile blend

**Table 2.** Emission rates and modeled intercellular concentrations of biogenic volatile organic compounds (BVOCs) in *Eugenia uniflora* sapling under filtered-air (FA) and ozone (FA + O₃) conditions.

| Class | BVOCs | Emission ($\mu\text{g gDM}^{-1} \text{h}^{-1}$) | | Intercellular ($\text{nmol mol}^{-1}$) | |
| --- | --- | --- | --- | --- | --- |
|  |  | FA | FA + O<sub>3</sub> | FA | FA + O<sub>3</sub> |
| ISO | Isoprene | 0.06 ± 0.02 | 0.07 ± 0.02 | 0.65 ± 0.24 | 0.57 ± 0.67 |
| MeSA | Methyl salicylate | 0.15 ± 0.07 | 0.18 ± 0.05 | 0.82 ± 0.29 | 0.73 ± 0.81 |
| MTs | 3-Carene | — | 0.08 ± 0.02 | — | 0.43 ± 0.47 |
|  | Terpinolene | — | 0.10 ± 0.03 | — | 0.55 ± 0.53 |
|  | β-Ocimene | — | 0.28 ± 0.17 | — | 2.32 ± 3.07 |
|  | β-Pinene | — | 0.10 ± 0.03 | — | 0.62 ± 0.78 |
|  | γ-Terpinene | — | 0.12 ± 0.03 | — | 0.71 ± 0.81 |
|  | **Total** |  | **0.68 ± 0.21** |  | **4.63 ± 5.56** |
| OSQTs | Curzerene | 0.48 ± 0.38 | 1.14 ± 1.14 | 2.36 ± 1.31 | 4.05 ± 5.59 |
|  | Germacrone | 0.58 ± 0.43 | 0.45 ± 0.16 | 3.24 ± 1.59 | 2.31 ± 2.48 |
|  | **Total** | **1.06 ± 0.66** | **1.59 ± 1.17** | **5.60 ± 1.83** | **6.35 ± 6.85** |
| SQTs | β-Guaiene | — | 0.08 ± 0.02 | — | 0.36 ± 0.38 |
|  | Caryophyllene | 0.42 ± 0.20 | — | 2.37 ± 0.90 | — |
|  | γ-Muurolene | 0.14 ± 0.07 | — | 0.86 ± 0.45 | — |
|  | Elixene | 1.02 ± 1.28 | 0.21 ± 0.09 | 4.76 ± 4.62 **a** | 1.01 ± 1.12 **b** |
|  | β-Elemene | 0.32 ± 0.36 | 0.39 ± 0.18 | 1.63 ± 1.53 | 1.83 ± 1.93 |
|  | γ-Elemene | 0.52 ± 0.60 **b** | 1.27 ± 0.74 **a** | 3.39 ± 3.31 | 5.42 ± 7.21 |
|  | γ-Gurjunene | 0.09 ± 0.04 | 0.09 ± 0.02 | 0.54 ± 0.17 | 0.40 ± 0.45 |
|  | **Total** | **2.52 ± 1.85** | **2.04 ± 0.88** | **13.54 ± 6.34** | **9.03 ± 10.52** |
| **Total BVOCs** |  | **3.79 ± 2.31** | **4.55 ± 1.61** | **20.60 ± 6.96** | **21.31 ± 22.41** |

Compounds are grouped by chemical class. Values are presented as mean ± standard deviation (n = 9 per treatment). Dashes (—) indicate compounds not detected. Bold values indicate class totals and total BVOC values. Class totals and total BVOCs represent the mean summed values for each chemical class and for all detected BVOCs, respectively. Differences between filtered-air (FA) control plants and ozone-exposed (FA + O<sub>3</sub>) plants were assessed within each row and measured variable using the Mann–Whitney U test. Different lowercase letters denote significant differences between treatments (p
< 0.05). ISO: isoprene; MeSA: methyl salicylate; MT: non-oxygenated monoterpene; OMT: oxygenated monoterpene; SQT: non-oxygenated sesquiterpene; OSQT: oxygenated sesquiterpene.

Despite these compound-specific changes, total BVOC emission rates and the summed emissions of SQTs and OSQTs remained unchanged under FA+O₃ treatment. Thus, O₃ primarily reorganized the composition of the emitted volatile blend through the induction of monoterpenes and the selective increase in γ-elemene, rather than increasing overall BVOC production.

Modeled intercellular BVOC concentrations were estimated from emission rates and gas-phase conductance parameters. Because these estimates were derived from the emission data, their qualitative composition broadly reflected the emitted profile. SQTs and OSQTs remained the dominant chemical class under both treatments, but summed emissions of these classes did not show significant difference as well as the total intercellular BVOC. However, compound-specific differences were observed. Elixene was the only BVOC whose modeled intercellular concentration decreased significantly under FA+O₃ treatment with large effect size, the other constitutive BVOCs did not show significant difference. Multivariate analyses confirmed that acute O₃ exposure induced a coordinated reorganization of BVOC metabolism. Emitted BVOC profiles were clearly separated according to treatment along the first principal component, which explained 48.7% of the total variance (Figure 2a). This separation was confirmed by PERMANOVA (R^2^ = 0.957; F =359; p = < 0.01). Multivariate dispersion did not differ significantly between treatments, as indicated by the betadisper test (F =0.71; p = 0.44), suggesting that the observed separation reflected differences in BVOC composition rather than unequal within-group variability. O₃-treated seedlings clustered with γ-elemene, curzerene, β-guaiene, and the induced monoterpenes, whereas filtered-air plants were primarily associated with elixene, caryophyllene, and γ-muurolene.

**Figure 2.**
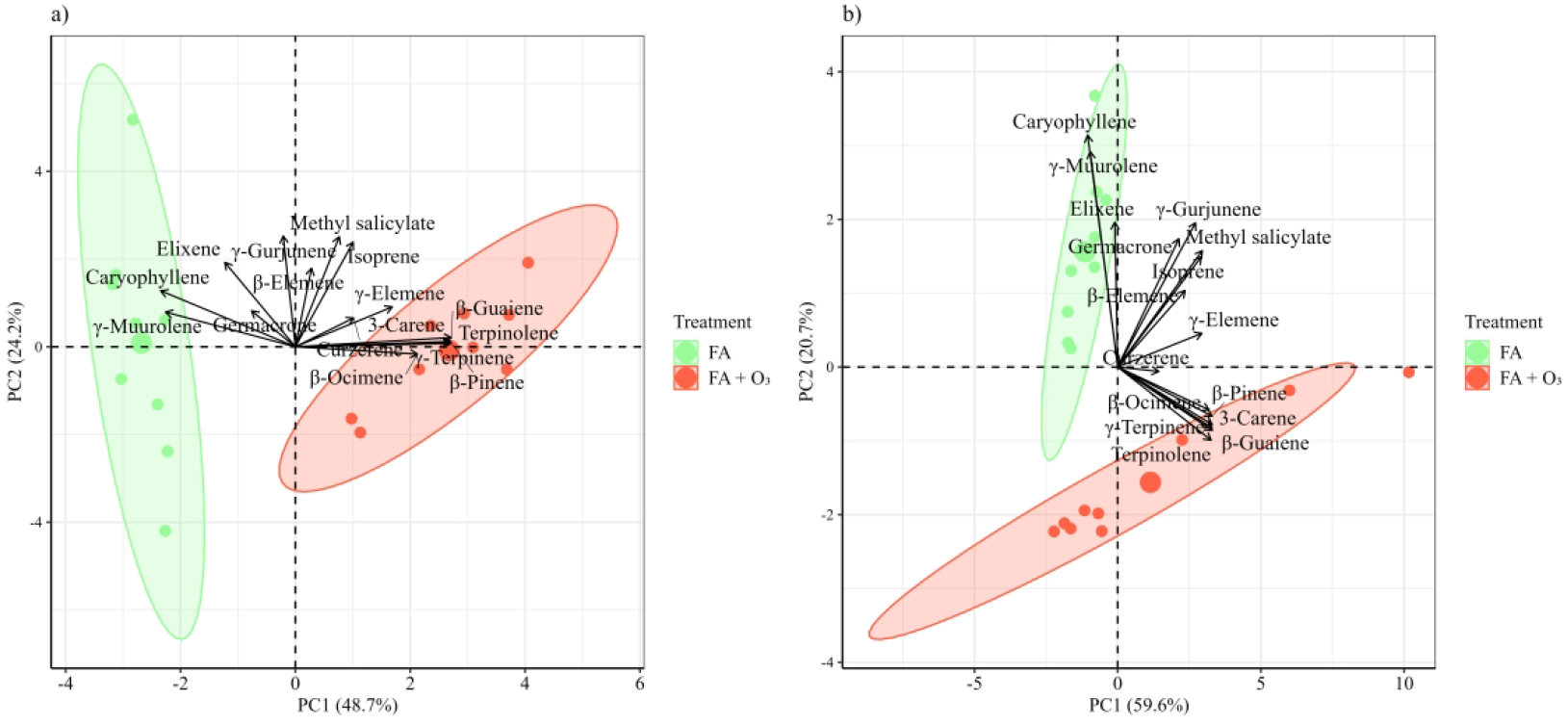
Principal component analysis (PCA) of (a) BVOC emission profiles and (b) modeled intercellular BVOC profiles of *Eugenia uniflora* L. saplings under control conditions (FA) and ozone exposure (FA + O₃). Points represent individual samples, while shaded ellipses indicate 95% confidence intervals for each treatment group. Vectors show the contribution of individual BVOCs to the ordination. Percentages on the axes indicate the variance explained by each principal component.

The modeled intercellular BVOC pool exhibited a similar but less pronounced response. Although O₃-treated plants tended to separate from filtered-air plants along the first principal component (Figure 2b), PERMANOVA (R^2^ = 0.968; F = 492; p = < 0.01) detected only a marginal treatment effect. Multivariate dispersion did not differ significantly between treatments (betadisper, F = 2.794; p = 0.130), indicating that the observed tendency was not attributable to unequal within-group variability. These results suggest that O₃ exerted a stronger influence on the composition of emitted BVOCs than on the overall composition of the intercellular volatile pool. Nevertheless, the same compounds largely explained treatment separation in both compartments, with O₃-treated plants associated with induced monoterpenes, β-guaiene, and γ-elemene, whereas filtered-air plants remained associated with elixene, germacrone, caryophyllene, and γ-muurolene.

Effect-size analysis further revealed that O₃ responses varied markedly among individual BVOCs and between emitted and modeled intercellular pools (Figure 3). Among emitted BVOCs (Figure 3a), γ-elemene exhibited the strongest positive effect and was the only compound significantly enhanced by O₃ exposure, consistent with its contribution to the compositional shift identified by the multivariate analyses. Moderate positive effect sizes were also observed for ISO, MeSA,curzerene and β-elemene, whereas elixene showed the strongest tendency toward higher emission under FA treatment; however, none of these responses was statistically significant (Table S4 available as Supplementary Data at Tree Physiology Online). In the modeled intercellular pool (Figure 3b), most compounds showed negative effect sizes, indicating a general tendency toward higher concentrations under FA treatment conditions. Elixene exhibited the strongest negative effect and was the only compound significantly reduced in the FA+ O_3_ treatment, whereas, curzerene, β-elemene, γ-elemene showed small positive effects.

**Figure 3.**
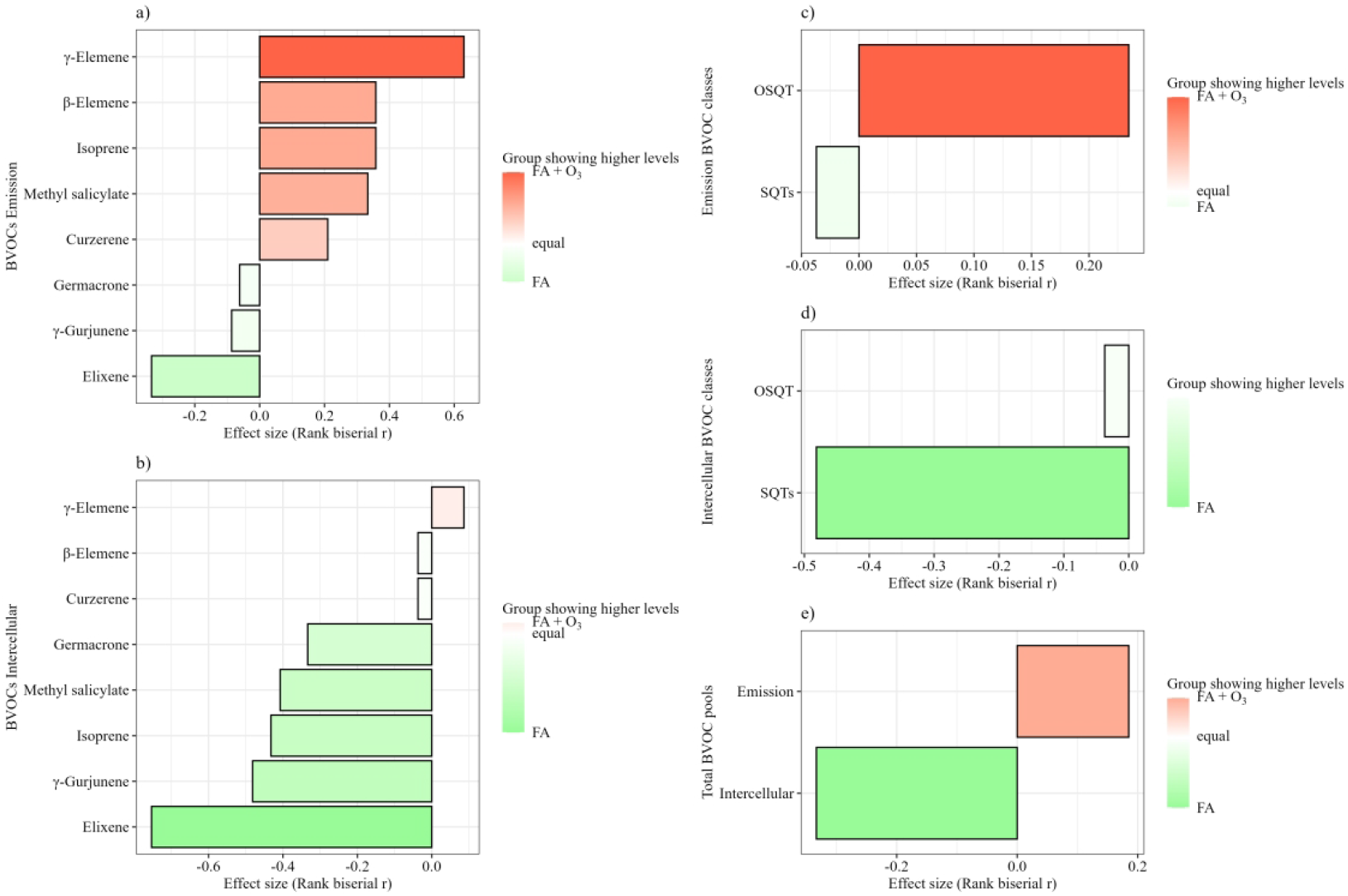
Effect sizes of O₃ exposure on emitted and modeled intercellular BVOC profiles in *Eugenia uniflora* L. seedlings. Rank-biserial correlation coefficients are shown for individual BVOCs in the emitted (a) and modeled intercellular (b) pools, for the predominant BVOC classes in the emitted (c) and modeled intercellular (d) pools, and for total BVOC pools (e). Positive values indicate higher levels in ozoe exposure (FA + O₃), whereas negative values indicate higher levels in filtered air (FA) plants. Bar colors indicate the treatment showing higher values. BVOC: Biogenic volatile organic compound.

OSQTs showed a positive effect size for emissions (Figure 3c), whereas SQTs showed a pronounced negative effect size in the modeled intercellular pool (Figure 3d); however, neither chemical classes differed significantly between treatments. Similarly, total emitted and modeled intercellular BVOC pools remained statistically unchanged, despite their opposite effect-size directions, with emissions tending to be higher under O₃ exposure and modeled intercellular concentrations tending to be higher under filtered-air conditions (Figure 3e).

Although no statistically significant changes were detected in total emitted and modeled intercellular BVOCs, the relative distribution of individual BVOCs between the modeled intercellular and emitted pools varied among compounds and was significantly altered under FA + O₃ (Figure 4 Table S6 available as Supplementary Data at Tree Physiology Online) Under FA treatment, ISO, germacrone, caryophyllene, γ-muurolene γ-elemene, were relatively more represented in the intercellular pool, whereas, curzerene and elixene, were more represented in emissions. Under FA+O₃ treatment, most induced monoterpenes, together with ISO and germacrone and elixene showed greater relative representation in the intercellular pool, while MeSA, curzerene and γ-elemene were relatively more represented in emissions

**Figure 4.**
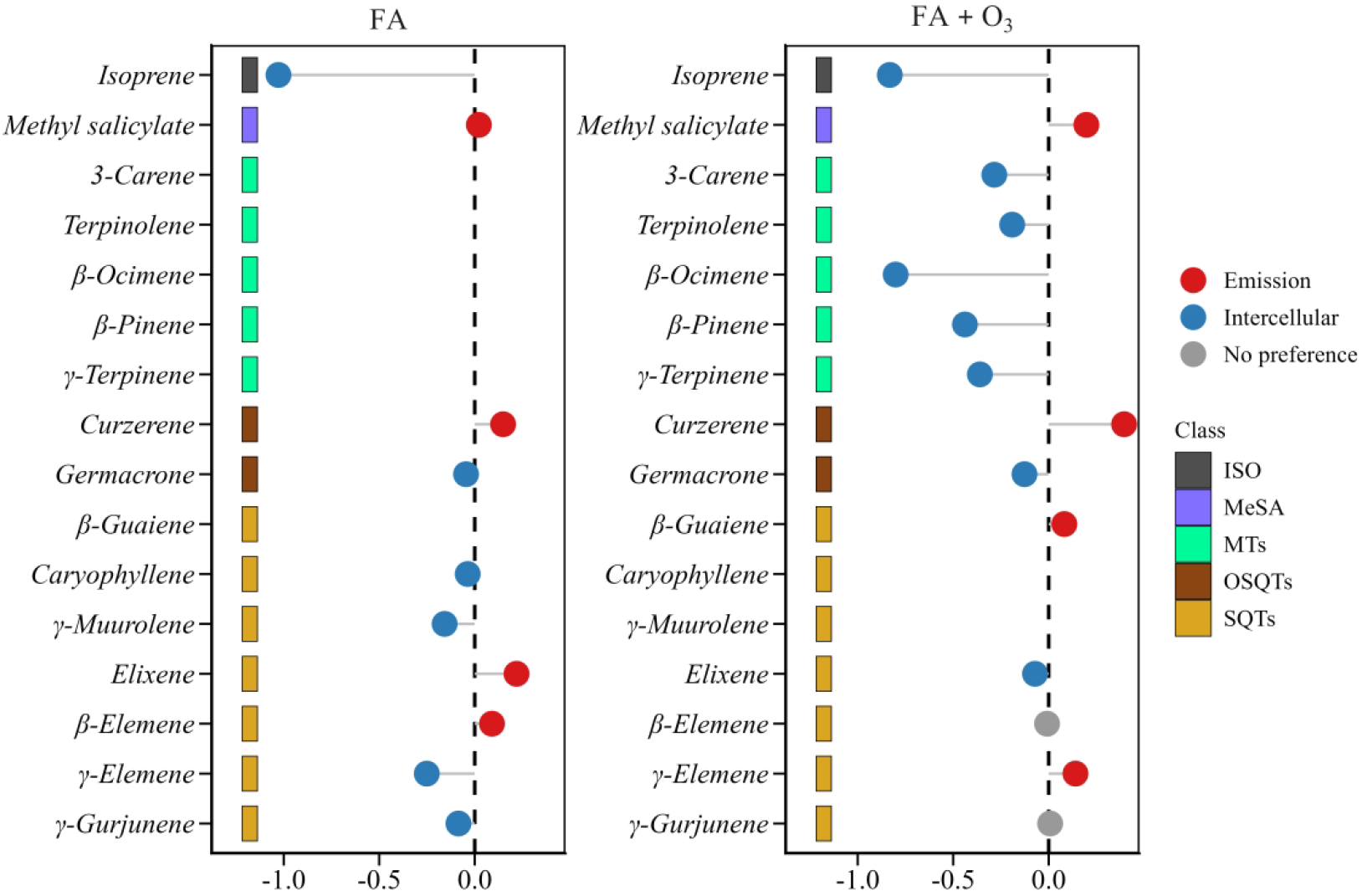
Emission-to-intercellular ratio of biogenic volatile organic compounds (BVOCs) in *Eugenia uniflora* under filtered air (FA) and ozone exposure (FA + O₃). Each point represents the log₂ ratio between emission and intercellular abundance for each BVOC. Positive values (red) indicate preferential partitioning toward emission, negative values (blue) indicate preferential accumulation in the intracellular compartment, and values close to zero (gray) indicate no preferential partitioning. Colored squares indicate the chemical class of each compound.

Although no statistically significant changes were detected in total emitted and modeled intercellular BVOCs, the relative distribution of individual BVOCs between the modeled intercellular and emitted pools varied among compounds and was significantly altered under FA + O₃ (Figure 4 Table S5-S6 available as Supplementary Data at Tree Physiology Online) Under FA treatment, ISO, germacrone, caryophyllene, γ-muurolene γ-elemene, were relatively more represented in the intercellular pool, whereas, curzerene and elixene, were more represented in emissions. Under FA+ O₃ treatment, most induced monoterpenes, together with ISO and germacrone and elixene showed greater relative representation in the intercellular pool, while MeSA, curzerene and γ-elemene were relatively more represented in emissions.

## Discussion

### Physiological and Redox Responses to Ozone

O₃ exposure did not impair photosynthetic performance under the conditions assessment. Net carbon assimilation increased significantly, whereas stomatal conductance, leaf water status, and chlorophyll concentrations remained unchanged. The maintenance of these physiological traits indicates that the applied O₃ regime did not exceed the capacity of *E. uniflora* to preserve photosynthetic function. A comparable maintenance of gas exchange and photosynthetic pigment concentrations was reported for *E. uniflora* saplings subjected to six days of O₃ exposure (AOT40 of 931 ppb h), despite the occurrence of localized foliar symptoms (Anselmo-Moreira et al. 2026). Similar transient stimulation of carbon assimilation has been reported under moderate or short-term O₃ exposure and has been interpreted as a hormetic response, whereby sublethal oxidative stress activates protective mechanisms without causing irreversible physiological damage (Calabrese et al. 2015; Agathokleous et al. 2019; Erofeeva 2022, 2023).

The preservation of photosynthetic performance was accompanied by distinct responses of the non-enzymatic antioxidant pools and a modest but significant decline in leaf pH. The AAt pool was already predominantly oxidized under FA and remained so under FA + O₃, suggesting limited capacity for further modulation of its redox state. The GSH pool exhibited the strongest response, with significant reductions in both total GSH and GSSG concentrations, while its redox status remained unchanged. The maintenance of GSH/GSSG status despite reductions in GSH suggests tight regulation of the redox status, potentially contributing to the preservation of cellular redox homeostasis during acute O₃ exposure (Gill and Tuteja 2010; Foyer and Noctor 2011; Noctor et al. 2012; Ding et al. 2020; Dorion et al. 2021; Ito and Ohkama-Ohtsu 2023; Noctor et al. 2024). Although bulk leaf pH cannot be directly interpreted as apoplastic pH, this response may be physiologically relevant because direct O₃ detoxification by apoplastic ascorbate is strongly pH-dependent. Thus, rather than eliciting a uniform antioxidant response, acute O₃ exposure appears to have differentially affected the two major non-enzymatic redox systems: the ascorbate pool remained predominantly oxidized, whereas glutathione maintained its redox status despite a contraction of the total pool. A comparable pattern was reported by Anselmo-Moreira et al. (2026) under a short-term O₃ regime with a lower accumulated AOT4, in which total glutathione showed a decreasing trend and GSH declined significantly, whereas the glutathione redox ratio remained unchanged. Engela et al. (2021) also reported reduced GSH concentrations in O₃-exposed *E. uniflora* under a prolonged exposure regime. Together, these studies suggest that alterations in the glutathione pool are a recurrent component of the response of *E. uniflora* to O_3_, while the contribution of the ascorbate system may depend on its initial redox state and the biochemical environment in which O₃ detoxification occurs.

In contrast, the total ascorbate pool remained unchanged. However, the lower proportion of reduced ascorbate together with the tendency toward higher DHA concentrations in O₃-treated plants suggests that ascorbate was more extensively oxidized during O₃ exposure. Because total ascorbate was maintained, these findings indicate increased turnover and regeneration of the ascorbate pool rather than depletion of antioxidant capacity. Such responses are compatible with activation of the ascorbate–glutathione cycle, in which continuous cycling between reduced and oxidized forms contributes to ROS detoxification under moderate oxidative stress (Bellini and De Tullio 2019; Ding et al. 2020; Xiao 2021). The concomitant reduction in carotenoid concentrations supports an increased demand for photoprotective antioxidants, reflecting their role in protecting the photosynthetic apparatus against oxidative damage (Strzałka et al. 2003; Nowroz et al. 2024). Thus, we believe that these findings indicate that maintenance of glutathione redox status, together with balance in the ascorbate pool and carotenoids, contributed to preserving cellular redox homeostasis during acute O₃ exposure.

Although the accumulated AOT40 exceeded the threshold generally considered harmful for vegetation, the relatively short exposure period may have favored acclimation rather than physiological impairment. This interpretation is according with Anselmo-Moreira et al. (2026), who observed physiological stability accompanied by localized foliar symptoms and antioxidant depletion after short-term O₃ exposure. Nevertheless, Engela et al. (2021) reported declines in photosynthetic performance and broader biochemical and metabolic alterations after 75 days of elevated O₃ exposure. These convergent and contrasting responses indicate that the outcome of O₃ exposure in *E. uniflora* depends not only on duration and cumulative dose, but also on the exposure system, environmental conditions, and timing of physiological and biochemical measurements. In the present study, the maintenance of photosynthetic function, together with selective antioxidant modulation, suggests that the plants retained sufficient physiological capacity to cope with acute ozone exposure.

### Enhanced Carbon Assimilation Supports Metabolic Reallocation

The stimulation of carbon assimilation observed under O₃ exposure was not accompanied by an increase in relative carbon loss through BVOC emissions, indicating that the additional carbon fixed during photosynthesis was not dissipated proportionally as volatile compounds. Instead, despite the pronounced restructuring of BVOC profiles, the proportion of recently assimilated carbon released to the atmosphere remained remarkably stable. This finding suggests that ozone exposure altered carbon partitioning rather than increasing the overall carbon investment in volatile emissions.

Terpene biosynthesis is energetically expensive and relies on a continuous supply of recently assimilated carbon (Loreto and Schnitzler 2010). Therefore, the enhanced carbon assimilation observed in *E. uniflora* may have provided the metabolic resources required to sustain O₃-induced synthesis of defensive terpenoids without increasing the proportional carbon loss through BVOC emission. This pattern is consistent with a reallocation of carbon resources toward defense while maintaining the relative cost of atmospheric volatile release (Yu and Blande, 2022; Shan and Jin 2025)

The comparison between emitted and modeled intercellular BVOC pools provided further insight into the fate of these compounds. Under O₃ exposure, several induced MTs showed greater intercellular accumulation than atmospheric release, indicating that their production was not necessarily followed by proportional emission. Their retention within the leaf may support local chemical protection while limiting carbon loss to the atmosphere (Niinemets et al. 2014; Yu and Blande 2022).

### Ozone-Induced changes in BVOC partitioning

Acute O₃ exposure substantially altered the BVOC profile of *E. uniflora*, with the response to oxidative stress expressed primarily through changes in composition and partitioning rather than through an increase in total BVOC emissions. This pattern is consistent with Anselmo-Moreira et al. (2026), who reported unchanged total BVOC emissions but significant alterations in the volatile profile of *E. uniflora* under short-term O₃ exposure. Together, these findings indicate that the response of this species to acute O₃ stress involves selective changes in individual BVOCs while maintaining relatively stable carbon loss through volatile emissions. In the present study, most constitutively detected compounds were maintained under FA, whereas several MTs, including 3-carene, β-cis-ocimene, γ-terpinene, terpinolene, and β-pinene, together with the SQT β-guaiene, appeared only in FA+ O₃ exposure. Conversely, γ-muurolene and caryophyllene were no longer detected in the ozone exposure. These qualitative changes indicate a selective reorganization of BVOC composition rather than a generalized stimulation of volatile emissions, consistent with O₃-induced changes in terpene profiles reported in other species (Loreto et al. 2004; Holopainen and Gershenzon 2010; Moura et al. 2022).

Among the constitutively detected compounds, γ-elemene exhibited the clearest positive response to O₃, with increased emission and a strong contribution to the O₃-associated shift in BVOC composition. This response suggests that γ-elemene may represent an important component of the acute response of *E. uniflora* to oxidative stress. SQT are highly reactive compounds that may contribute to plant responses to atmospheric oxidants through several non-mutually exclusive mechanisms, including reactions with ROS and O₃ and modulation of oxidative signaling (Holopainen and Gershenzon 2010; Yu and Blande 2022; Costa et al. 2025). The appearance of several MTs exclusively under FA+O₃ indicates that the response extended beyond changes in constitutive SQT emissions and involved a broader reorganization of terpene composition. Because many MTs and SQTs react rapidly with O₃, their potential protective effects may occur both within the leaf and after release to the atmosphere (Fares et al. 2007; Ghimire et al. 2017; Mochizuki et al. 2017; Yu and Blande 2022). However, the contribution of emitted BVOCs to O₃ removal near the leaf surface should be interpreted cautiously, as its relevance depends on compound-specific reactivity, emission rates, atmospheric mixing, and the spatial proximity between BVOC release and O₃ uptake.

Importantly, the simultaneous assessment of emitted BVOCs and modeled intercellular concentrations provides an additional perspective on BVOC partitioning that cannot be obtained from emission measurements alone. Our previous study suggested a potential decoupling between SQT production and atmospheric release under O₃ exposure, as relatively stable or enhanced internal terpenoid pools occurred without a corresponding increase in emissions, raising the possibility of altered retention, allocation, or in situ consumption within the leaf (Anselmo-Moreira et al. 2026). The present results extend this interpretation by showing that O₃ modified the relative distribution of individual BVOCs between the modeled intercellular and emitted pools. Thus, changes in emission rates cannot necessarily be interpreted as direct proxies for changes in BVOC production. A compound may be synthesized but preferentially retained within the leaf, released to the atmosphere, or consumed through chemical reactions before emission. This distinction is particularly relevant under O₃ stress because highly reactive BVOCs present in the intercellular air spaces may encounter O₃ entering through the stomata and potentially contribute to an internal chemical barrier against oxidative stress (Yu and Blande 2022). The compound-specific responses observed support changes in BVOC partitioning under ozone stress. Under FA + O₃, several induced MTs showed greater relative representation in the modeled intercellular pool, whereas MeSa, curzerene, γ-elemene were relatively more represented in emissions. O₃ therefore did not induce a uniform shift toward either intercellular accumulation or atmospheric release but instead altered BVOC partitioning in a compound-specific manner. These differences might reflect variation in physicochemical properties, storage capacity, gas-phase conductance, and chemical reactivity within the leaf. Thus, the O₃-induced BVOC response of *E. uniflora* appears to involve compound-specific changes in partitioning, reflecting a balance among intercellular accumulation, potential chemical consumption, and atmospheric release rather than a simple increase or decrease in total emissions.

## Conclusion

O₃ exposure induced an integrated physiological response in *E. uniflora*, characterized by enhanced carbon assimilation, maintenance of glutathione redox status, and compound-specific changes in BVOC partitioning. Increased carbon assimilation may have supported carbon availability for stress-related processes, while stable total BVOC emissions prevented additional carbon loss through volatile release. At the same time, changes in modeled intercellular BVOC concentrations indicate that part of the volatile response was retained within the leaf rather than expressed as increased atmospheric emission. These findings show that BVOC production, intercellular accumulation, potential chemical consumption, and emission are interconnected processes that collectively determine the fate of volatile carbon under oxidative stress. Thus, the response of *E. uniflora* to acute O₃ exposure was characterized not by increased total BVOC emission, but by a reorganization of BVOC composition and partitioning between the modeled intercellular and emitted pools, while maintaining photosynthetic performance, glutathione redox status, and stable carbon loss through volatile emissions. Beyond advancing our mechanistic understanding of plant responses to O₃, this study provides a physiological framework for future comparative assessments of tree species used in urban greening. Integrating traits related to carbon acquisition, redox homeostasis, and BVOC partitioning may contribute to the identification of species better suited to maintain ecosystem services in O₃-polluted metropolitan environments.

## Acknoledgements

The authors would like to thank the Fundação de Pesquisa do Agronegócio (FUNDEPAG) for its support (grant 2022/039228). AN acknowledge financial support and the scholarships from CAPES (Coordenação de Aperfeiçoamento de Pessoal de Nível Superior). FAM and BRBC are grateful for their postdoctoral grants from FAPESP (grants 2022/07326-8, 2022/13213-1, and 2022/11143-6, respectively). CMF acknowledges CNPq (National Council for Scientific and Technological Development) for the Research Productivity Grant (302188/2022-3). SRS acknowledges FUNDEPAG for the Research Grant (2022/2672).

## Author’sContribution

AN: Investigation; Data curation; Formal analysis; Writing – original draft; Writing – review & editing. FAM and BRBC: Investigation; Data curation; Writing – original draft; Writing review & editing. MCS: Investigation; Data curation. CMF: Investigation; Writing – review & editing. SRS: Conceptualization; Funding acquisition; Project administration; Resources; Supervision; Writing original draft; Writing – review & editing.

## Supplementary Data

Supplementary Data are available at *Tree Physiology* Online and include additional datasets, Supplementary Figureures S1–S5, and Supplementary Tables S1–S4

## Funding

This work was supported by grants from the São Paulo Research Foundation (FAPESP), Fundação de Desenvolvimento da Pesquisa do Agronegócio (FUNDEPAG), Coordination for the Improvement of Higher Education Personnel (CAPES), and the Brazilian National Council for Scientific and Technological Development (CNPq)

## Conflict of interest

The authors declare that there are no conflicts of interest.

## Data Availability

All data are incorporated into the article and its online supplementary material

## Notes

### Competing Interest Statement

The authors have declared no competing interest.

## References

Agathokleous E, Araminiene V, Belz RG, Calatayud V, De Marco A, Domingos M, Feng Z, Hoshika Y, Kitao M, Koike T, et al. 2019. A quantitative assessment of hormetic responses of plants to ozone. Environ Res. 176:108527. 10.1016/j.envres.2019.108527.

Ainsworth EA, Yendrek CR, Sitch S, Collins WJ, Emberson LD. 2012. The effects of tropospheric ozone on net primary productivity and implications for climate change. Annu Rev Plant Biol. 63:637–661. 10.1146/annurev-arplant-042110-103829.

Anselmo-Moreira F, Claude A, Nascimento A, Costa BRB, Hurtado-Caceres I, Rocco M, Staudt M, Fornaro A, Borbon A, Furlan CM, Souza SR. 2026. Drought modulates ozone stress through BVOCs, antioxidant defenses, and metabolic responses in a tropical tree. Planta. 10.1007/s00425-026-05033-8

Anselmo-Moreira F, da Silva Pedrosa G, da Silva IL, do Nascimento A, dos Santos TC, Catharino ELM, Gomes EPC, Borbon A, Fornaro A, de Souza SR. 2025. Biogenic volatile organic compound (BVOC) emission profiles from native Atlantic Forest trees: seasonal variation and atmospheric implications in southeastern Brazil. Urban For Urban Green. 104:128645. 10.1016/j.ufug.2024.128645.

Bellini E, De Tullio MC. 2019. Ascorbic acid and ozone: novel perspectives to explain an elusive relationship. Plants. 8(5):122. 10.3390/plants8050122.

Borbon A, Fornaro A, Oliveira AP, Souza SR, Brito JF, Jaffrezo J-L, Staudt M, Ynoue RY, Codato G, et al. 2026. The BIOMASP+ project on biosphere–atmosphere exchanges and their role in air pollution in the subtropical megacity of São Paulo: motivations, methods, and preliminary observations. Bull Am Meteorol Soc. 107(4):E902–E921. 10.1175/BAMS-D-23-0161.1.

Burrows FJ, Milthorpe FL. 1976. Stomatal conductance in the control of gas exchange. In: Kozlowski TT, editor. Soil Water Measurement, Plant Responses, and Breeding for Drought Resistance. New York (NY): Academic Press. Chapter 3; p. 103–152.

Calabrese EJ, Dhawan G, Kapoor R, Iavicoli I, Calabrese V. 2015. What is hormesis and its relevance to healthy aging and longevity? Biogerontology. 16(6):693–707. 10.1007/s10522-015-9601-0

Cheesman AW, Brown F, Artaxo P, Farha MN, Folberth GA, Hayes FJ, Heinrich VHA, Hill TC, Mercado LM, Oliver RJ, et al. 2024. Reduced productivity and carbon drawdown of tropical forests from ground-level ozone exposure. Nat Geosci. 17(10):1003–1007. 10.1038/s41561-024-01530-1.

Cohen J. 1988. Statistical power analysis for the behavioral sciences. 2nd ed. Hillsdale(NJ): Lawrence Erlbaum Associates.

Costa BRB, Anselmo-Moreira F, Nascimento A, Pedrosa GS, Catharino ELM, Borbon A, Fornaro A, Furlan CM, Souza SR. 2025. Unveiling sesquiterpene emissions in dominant trees of a Brazilian Atlantic Forest remnant. Atmos Environ X. 27:100358. 10.1016/j.aeaoa.2025.100358.

Costa JS, Barroso AS, Mourão RHV, da Silva JKR, Maia JGS, Figureueiredo PLB. 2020. Seasonal and antioxidant evaluation of essential oil from *Eugenia uniflora* L., curzerene-rich, thermally produced in situ. Biomolecules. 10(2):328. 10.3390/biom10020328.

Ding H, Wang B, Han Y, Li S. 2020. The pivotal function of dehydroascorbate reductase in glutathione homeostasis in plants. Journal of Experimental Botany. 71(12):3405–3416. 10.1093/jxb/eraa107

Dorion S, Ouellet JC, Rivoal J. 2021. Glutathione metabolism in plants under stress: beyond reactive oxygen species detoxification. Metabolites. 11(9):641. 10.3390/metabo11090641.

Engela MRG da S, Furlan CM, Esposito MP, Fernandes FF, Carrari E, Domingos M, Paoletti E, Hoshika Y. 2021. Metabolic and physiological alterations indicate that the tropical broadleaf tree *Eugenia uniflora* L. is sensitive to ozone. Sci Total Environ. 769:145080. 10.1016/j.scitotenv.2021.145080.

Erofeeva EA. 2022. Hormesis in plants: its common occurrence across stresses. Curr Opin Toxicol. 30:100333. 10.1016/j.cotox.2022.02.006.

Erofeeva EA. 2023. Hormetic effects of abiotic environmental stressors in woody plants in the context of climate change. J For Res. 34:7–19. 10.1007/s11676-022-01547-0.

Fall R, Monson RK. 1992. Isoprene emission rate and intercellular isoprene concentration as influenced by stomatal distribution and conductance. Plant Physiol. 100(2):987–992. 10.1104/pp.100.2.987.

Fares S, Loreto F, Kleist E, Wildt J. 2007. Stomatal uptake and stomatal deposition of ozone in isoprene and monoterpene emitting plants. Plant Biol. 9:e69–e78. 10.1111/j.1438-8677.2007.00062.x.

Filzmoser P, Hron K, Templ M. 2018. Applied compositional data analysis: with worked examples in R. Cham (Switzerland): Springer International Publishing. 10.1007/978-3-319-96422-5.

Foyer CH, Noctor G. 2011. Ascorbate and glutathione: the heart of the redox hub. Plant Physiol. 155(1):2–18. 10.1104/pp.110.167569.

Ghimire RP, Kivimäenpää M, Kasurinen A, Häikiö E, Holopainen T, Holopainen JK. 2017. Herbivore-induced BVOC emissions of Scots pine under warming, elevated ozone and increased nitrogen availability in an open-field exposure. Agric For Meteorol. 242:21–32. 10.1016/j.agrformet.2017.04.008.

Gill SS, Tuteja N. 2010. Reactive oxygen species and antioxidant machinery in abiotic stress tolerance in crop plants. Plant Physiol Biochem. 48(12):909–930. 10.1016/j.plaphy.2010.08.016.

González L, González-Vilar M. 2001. Determination of relative water content. In: Reigosa Roger MJ, editor. Handbook of Plant Ecophysiology Techniques. Dordrecht: Springer. p. 207–212. 10.1007/0-306-48057-3_14.

Guenther AB, Jiang X, Heald CL, Sakulyanontvittaya T, Duhl T, Emmons LK, Wang X. 2012. The Model of Emissions of Gases and Aerosols from Nature version 2.1 (MEGAN2.1): an extended and updated framework for modeling biogenic emissions. Geosci Model Dev. 5(6):1471–1492. 10.5194/gmd-5-1471-2012.

Holopainen JK, Gershenzon J. 2010. Multiple stress factors and the emission of plant VOCs. Trends Plant Sci. 15(3):176–184. 10.1016/j.tplants.2010.01.006.

Ito T, Ohkama-Ohtsu N. 2023. Degradation of glutathione and glutathione conjugates in plants. Journal of Experimental Botany. 74(11):3313–3327. 10.1093/jxb/erad018.

Karl T, Fall R, Rosenstiel TN, Prazeller P, Larsen B, Seufert G, Lindinger W. 2002. On-line analysis of the 13CO₂ labeling of leaf isoprene suggests multiple subcellular origins of isoprene precursors. Planta. 215(6):894–905. 10.1007/s00425-002-0825-2.

Kassambara A, Mundt F. 2020. factoextra: extract and visualize the results of multivariate data analyses. R package version 1.0.7. https://CRAN.R-project.org/package=factoextra.

Lê S, Josse J, Husson F. 2008. FactoMineR: an R package for multivariate analysis. Journal of Statistical Software. 25(1):1–18. 10.18637/jss.v025.i01.

Lichtenthaler HK. 1987. Chlorophylls and carotenoids: pigments of photosynthetic biomembranes. In: Douce R, Packer L, editors. Methods Enzymol. 148:350–382. 10.1016/0076-6879(87)48036-1.

Liu S, Chen J, Han W. 2022. Comparison of pretreatment, preservation and determination methods for foliar pH of plant samples. J Plant Ecol. 15(4):673–682. 10.1093/jpe/rtac012.

LI-COR Inc. 2002. LI-6400 portable photosynthesis system: instruction manual. Lincoln (NE): LI-COR Biosciences.

López A, Montaño A, García P, Garrido A. 2005. Quantification of ascorbic acid and dehydroascorbic acid in fresh olives and in commercial presentations of table olives. Food Sci Technol Int. 11(3):199–204. 10.1177/1082013205054421.

Loreto F, Pinelli P, Manes F, Kollist H. 2004. Impact of ozone on monoterpene emissions and evidence for an isoprene-like antioxidant action of monoterpenes emitted by *Quercus ilex* leaves. Tree Physiol. 24(4):361–367. 10.1093/treephys/24.4.361.

Loreto F, Schnitzler J-P. 2010. Abiotic stresses and induced BVOCs. Trends Plant Sci. 15(3):154–166. 10.1016/j.tplants.2009.12.006.

Mills G, Pleijel H, Malley CS, Sinha B, Cooper OR, Schultz MG, Neufeld HS, Simpson D, Sharps K, Feng Z, Gerosa G, Harmens H, Kobayashi K, Saxena P, Paoletti E, Sinha V, Xu X. 2018. Tropospheric Ozone Assessment Report: present-day tropospheric ozone distribution and trends relevant to vegetation. Elementa (Wash D C). 6:47. 10.1525/elementa.302.

Minocha R, Martinez G, Lyons B, Long S. 2009. Development of a standardized methodology for quantifying total chlorophyll and carotenoids from foliage of hardwood and conifer tree species. Can J For Res. 39(4):849–861. 10.1139/X09-015.

Mochizuki T, Watanabe M, Koike T, Tani A. 2017. Monoterpene emissions from needles of hybrid larch F1 (*Larix gmelinii* var. *japonica* × *Larix kaempferi*) grown under elevated carbon dioxide and ozone. Atmos Environ. 148:197–202. 10.1016/j.atmosenv.2016.10.041.

Monks PS, Archibald AT, Colette A, Cooper O, Coyle M, Derwent R, Fowler D, Granier C, Law KS, Mills GE, Stevenson DS, Tarasova O, Thouret V, von Schneidemesser E, Sommariva R, Wild O, Williams ML. 2015. Tropospheric ozone and its precursors from the urban to the global scale: from air quality to short-lived climate forcer. Atmos Chem Phys. 15:8889–8973. 10.5194/acp-15-8889-2015

Moura BB, Bolsoni VP, de Paula MD, Dias GM, de Souza SR. 2022. Ozone impact on emission of biogenic volatile organic compounds in three tropical tree species from the Atlantic Forest remnants in Southeast Brazil. Front Plant Sci. 13:879039. 10.3389/fpls.2022.879039.

Niinemets Ü, Fares S, Harley P, Jardine KJ. 2014. Bidirectional exchange of biogenic volatiles with vegetation: emission sources, reactions, breakdown and deposition. Plant Cell Environ. 37(8):1790–1809. 10.1111/pce.12322.

Niinemets Ü, Reichstein M, Staudt M, Seufert G, Tenhunen JD. 2002. Stomatal constraints may affect emission of oxygenated monoterpenoids from the foliage of *Pinus pinea*. Plant Physiol. 130(3):1371–1385. 10.1104/pp.009670.

Noctor G, Cohen M, Trémulot L, Châtel-Innocenti G, Van Breusegem F, Mhamdi A. 2024. Glutathione: a key modulator of plant defence and metabolism through multiple mechanisms. J Exp Bot. 75(15):4549–4572. 10.1093/jxb/erae194.

Noctor G, Mhamdi A, Chaouch S, Han Y, Neukermans J, Márquez-García B, Queval G, Foyer CH. 2012. Glutathione in plants: an integrated overview. Plant Cell Environ. 35:454–484. 10.1111/j.1365-3040.2011.02400.x

Nowroz F, Hasanuzzaman M, Siddika A, Parvin K, Caparros PG, Nahar K, Prasad PVV. 2024. Elevated tropospheric ozone and crop production: potential negative effects and plant defense mechanisms. Frontiers in Plant Science. 14:1244515. 10.3389/fpls.2023.1244515.

Peñuelas J, Staudt M. 2010. BVOCs and global change. Trends Plant Sci. 15(3):133–144. 10.1016/j.tplants.2009.12.005.

Pérez-Harguindeguy N, Díaz S, Garnier E, Lavorel S, Poorter H, Jaureguiberry P, Bret-Harte MS, Cornwell WK, Craine JM, et al. 2013. New handbook for standardized measurement of plant functional traits worldwide. Aust J Bot. 61(3):167–234. 10.1071/BT12225.

R Core Team. 2024. R: a language and environment for statistical computing. Vienna (Austria): R Foundation for Statistical Computing. https://www.r-project.org/.

Sala-Carvalho WR, Montessi-Amaral FP, Esposito MP, Campestrini R, Rossi M, Peralta DF, Furlan CM. 2022. Metabolome of *Ceratodon purpureus* (Hedw.) Brid., a cosmopolitan moss: the influence of seasonality. Planta. 255(4):77. 10.1007/s00425-022-03857-8.

Shan Y, Jin S. 2025. Biosynthetic machinery to abiotic stress-driven emission: decoding multilayer regulation of volatile terpenoids in plants. Antioxidants (Basel). 14(6):673. 10.3390/antiox14060673.

Singh AA, Ghosh A, Agrawal M, Agrawal SB. 2023. Secondary metabolites responses of plants exposed to ozone: an update. Environ Sci Pollut Res. 30(38):88281–88312. 10.1007/s11356-023-28634-2.

Souza SR, Pagliuso JD. 2009. Design and assembly of an experimental laboratory for the study of atmosphere–plant interactions in the system of fumigation chambers. Environ Monit Assess. 158(1):243–249. 10.1007/s10661-008-0578-x.

Stekhoven DJ, Bühlmann P. 2012. MissForest—non-parametric missing value imputation for mixed-type data. Bioinformatics. 28(1):112–118. 10.1093/bioinformatics/btr597.

Strzałka K, Kostecka-Gugała A, Latowski D. 2003. Carotenoids and environmental stress in plants: significance of carotenoid-mediated modulation of membrane physical properties. Russ J Plant Physiol. 50(2):168–173. 10.1023/A:1022960828050.

Vainonen JP, Kangasjärvi J. 2015. Plant signalling in acute ozone exposure. Plant Cell Environ. 38(2):240–252. 10.1111/pce.12273.

Vickers CE, Gershenzon J, Lerdau MT, Loreto F. 2009. A unified mechanism of action for volatile isoprenoids in plant abiotic stress. Nat Chem Biol. 5(5):283–291. 10.1038/nchembio.158.

von Caemmerer S, Farquhar GD. 1981. Some relationships between the biochemistry of photosynthesis and the gas exchange of leaves. Planta. 153:376–387. 10.1007/BF00384257.

Weatherley PE. 1950. Studies in the water relations of the cotton plant: I. The field measurement of water deficits in leaves. New Phytol. 49(1):81–97. 10.1111/j.1469-8137.1950.tb05146.x.

Wedow JM, Ainsworth EA, Li S. 2021. Plant biochemistry influences tropospheric ozone formation, destruction, deposition, and response. Trends Biochem Sci. 46(12):992–1002. 10.1016/j.tibs.2021.06.007.

Xiao M, Li Z, Zhu L, Wang J, Zhang B, Zheng F, Zhao B, Zhang H, Wang Y, Zhang Z. 2021. The multiple roles of ascorbate in the abiotic stress response of plants: antioxidant, cofactor, and regulator. Front Plant Sci. 12:598173. 10.3389/fpls.2021.598173.

Yu H, Blande JD. 2021. Diurnal variation in BVOC emission and CO₂ gas exchange from above- and belowground parts of two coniferous species and their responses to elevated O₃. Environ Pollut. 278:116830. 10.1016/j.envpol.2021.116830.

Yu H, Blande JD. 2022. A potential ozone defense in intercellular air space: clues from intercellular BVOC concentrations and stomatal conductance. Sci Total Environ. 852:158456. 10.1016/j.scitotenv.2022.158456.

Zuo Z, Weraduwage SM, Huang T, Sharkey TD. 2025. How volatile isoprenoids improve plant thermotolerance. Trends Plant Sci. 30(11):1237. 10.1016/j.tplants.2025.05.004.

